# Selective enhancement of fronto-temporal mechanisms for adaptive memory

**DOI:** 10.64898/2025.12.19.695543

**Authors:** Benjamin Slater, Timothy Griffiths, Hugo Caffaratti, Jérôme Sallet, Patrick Degenaar, Marcus Kaiser, Alexander Easton, Yukiko Kikuchi, Christopher Petkov

**Author notes:** Senior author (Christopher Petkov).

## Abstract

There is substantial scientific interest in advancing knowledge on the control mechanisms supporting adaptive memory and improving approaches that can enhance cognition through brain stimulation. We implemented a non-invasive focal Transcranial Ultrasound Stimulation (TUS) approach with known longer-lasting post-stimulation effects in two rhesus macaques performing a novel context-dependent memory-sequencing task implemented on multiple touchscreens within their home units. Consistently in both monkeys, TUS to the anterior, but not posterior, medial temporal lobe enhanced performance under stable memory-sequencing contexts. TUS to the medial prefrontal cortex, on the other hand, selectively improved performance when contexts were unstable and the monkey needed to adapt to both a change in context and temporal sequence. These findings provide positive causal evidence of a functional dissociation within fronto-temporal memory circuits and pave the way for advancing non-invasive approaches to improve cognitive function and the study of neural circuits under focal enhancement across species.

**IMPACT STATEMENT:** Transcranial ultrasound stimulation of the prefrontal-hippocampal network selectively enhances context-guided memory in macaques, demonstrating that non-invasive brain stimulation can causally improve, rather than only disrupt, memory circuits.

## INTRODUCTION

Adaptive behaviour is dependent on the brain resolving a central computational challenge: balancing stable memory retrieval based on successful prior experience, with flexibility when environmental contingencies require a change in behaviour. Such cognitive functions are thought to depend on dynamic interactions between prefrontal and medial temporal lobe (MTL) nodes and there is substantial interest in how the functions of this system could potentially be selectively causally disrupted or enhanced with brain stimulation (Polanía et al., 2018). Here, we aimed to dissect the functional roles of the anterior and posterior MTL, and the medial prefrontal cortex (mPFC) using a promising non-invasive brain stimulation approach with spatial specificity, low-intensity Transcranial Ultrasound Stimulation (TUS), implemented with two rhesus macaque monkeys as animal models of complex cognition.

Fronto-temporal interactions are well established as critical for memory and flexible behaviour (Barker et al., 2017; Barker and Warburton, 2020; Cernotova et al., 2021; Easton and Gaffan, 2002; Gaffan et al., 2002; Malik et al., 2022; Weilbächer and Gluth, 2016; Wirt and Hyman, 2017). Anterior hippocampal regions support context-dependent episodic memory (Liang et al., 2013), posterior hippocampal regions encode fine-grained spatial and temporal detail (Komorowski et al., 2009), and the PFC resolves competing associations and updates contingencies when environmental conditions change (Eichenbaum and Howard, 2017; Preston and Eichenbaum, 2013; Rich and Shapiro, 2007; Simons and Spiers, 2003). Rodent studies show that disrupting hippocampal-frontal communication impairs encoding but not retrieval (Spellman et al., 2015), extending lesion-based findings from humans and nonhuman animal models (Euston et al., 2012). Progress in understanding these circuits depends on approaches that can focally perturb deep brain structures and enable causal dissection of pathway-specific functions to reveal conditions under which stimulation may enhance, rather than simply disrupt, cognitive function.

Low-intensity TUS has emerged as a promising approach for such causal dissection. Its capability to pass focused sonic pulses through the skull enables millimetre-scale targeting of structures that are inaccessible to other non-invasive methods for neuromodulation (Blackmore et al., 2023; Yoo et al., 2011). Research with animal models has further established ‘offline’ stimulation protocols whose effects outlast the relatively brief period of sonication by tens of minutes, if not hours (Atkinson-Clement et al., 2025; Bault et al., 2024). Although TUS can modulate behaviour and neural activity in either direction, its functional consequences remain difficult to predict: a recent meta-analysis and resource on TUS studies in healthy human participants highlights substantial variability in the directionality and magnitude of reported effects (Caffaratti et al., 2025). This underscores the need for mechanistic studies that link stimulation site, task demands, and behavioural outcome.

Research in nonhuman animals, including nonhuman primates, has been essential for developing TUS as a translational neuromodulation tool for therapeutic use in humans (Atkinson-Clement et al., 2025; Bongioanni et al., 2021; Folloni et al., 2021, 2019; Fouragnan et al., 2019; Khalighinejad et al., 2020; Verhagen et al., 2019). Yet most pioneering primate studies to date have reported disruption rather than enhancement. For example, sonication of anterior prefrontal regions in macaques during value-based decision making reduced the integration of reward probability and magnitude (Bongioanni et al., 2021). Similarly, sonicating anterior cingulate cortex (ACC) activity with TUS impaired macaques’ use of counterfactual choice-value information to guide behavioural change (Fouragnan et al., 2019). Using an offline TUS protocol closely matching the one used here, TUS of the ACC has also been shown to slow learning and prolong information sampling in macaques under high motivational and cognitive demand (Boroujeni et al., 2022). What is surprising, is that the general TUS ‘offline’ stimulation protocol implemented in these primate studies, and in the current one (e.g., 5-10 Hz stimulation frequency, 10% duty cycle), have been shown to be excitatory in rodents both in its immediate effects on neuronal responses (Murphy et al., 2024) and its ‘offline’ effects after the sonication period (Kim et al., 2024). Therefore, although prior nonhuman primate studies demonstrate that TUS can causally disrupt frontal computations, the question remains as to whether such protocols always produce disruption, or whether they can also improve cognitive function in a site- and function-specific manner.

To advance insights on the impact of TUS on cognitive function, we tested how focal low-intensity TUS applied to the MTL, including the hippocampus, or to the medial PFC causally alters context-guided memory sequencing in rhesus macaques. We developed a novel free-moving, home-unit touchscreen task that required learning and recalling object sequences under two different contextual scenarios. Context was conveyed by the background colour on the touch screens that either remained stable across trials on the screens or shifted unpredictably within or across trials, allowing us to dissociate memory processes favouring stability versus flexibility. Across several weeks, each monkey completed a within-subjects, counterbalanced series of sessions in which we applied an ‘offline’ low-intensity TUS protocol to the medial PFC (mPFC), anterior MTL (aMTL), or posterior MTL (pMTL), comparing performance in each condition to interleaved sham sessions. Behavioural effects after TUS or sham were quantified using logistic regression models to capture fine-grained modulation of performance patterns across stimulation sites and context conditions.

This context-guided memory sequencing task design builds on a broader literature examining context- and temporal-order learning in rodents and primates, including work on ordinal-transfer errors in context-guided sequence tasks (Reeders et al., 2021), temporal order learning in monkeys (Templer and Hampton, 2013), and neural dynamics during touchscreen context-guided behaviour (Abbaspoor and Hoffman, 2024). A more technical description of the task is a context-dependent biconditional paired-associate task with a temporal-order component in visual sequences.

Our TUS stimulation parameters were chosen to fall within a parameter range considered more likely to enhance, rather than disrupt, local neural activity (Caffaratti et al., 2025; Kim et al., 2024; Murphy et al., 2024). However, given the limited predictability of the directionality of TUS effects, we evaluated both disruption and enhancement hypotheses. If TUS primarily induced transient disruption, we expected stimulation of the aMTL to impair performance under stable contextual conditions. Behaviour in this context relies on retrieving generalisable relational structure across sequences, an operation linked to anterior hippocampal involvement in context-dependent episodic memory and schema-like integration (Nadel et al., 2013; Ryan et al., 2010). Disrupting this pathway should therefore compromise the use of stable contextual cues. We likewise predicted that disrupting mPFC function would impair performance under unstable contextual conditions, where flexible updating, conflict resolution, and adaptation to shifting contingencies are required (Benchenane et al., 2010; Place et al., 2016).

Conversely, if stimulation acted in an enhancing manner, the opposite pattern should emerge. Enhancing aMTL function would be expected to facilitate performance under stable conditions, where strengthening access to generalisable relational representations could improve sequence recall. Enhancing mPFC function should benefit performance in unstable contexts, where improved interference resolution and more efficient updating of context-sequence associations would directly support behavioural flexibility.

All procedures adhered to ethical principles for nonhuman primate neuroscience, including minimising animal numbers under the 3Rs framework (Hubrecht and Carter, 2019; Petkov et al., 2022). Following this principle, testing was limited to two animals, each serving as their own within-subjects control across all stimulation and sham conditions. We report only those effects that were consistent in direction and statistically reliable in both animals, irrespective of differences in training history.

## RESULTS

The present study conducted a context-guided sequence learning task using multiple touchscreens within the animals’ home environment, as we evaluated the impact on their behavioural performance by applying transcranial ultrasound stimulation (TUS) or sham stimulation to the medial prefrontal cortex and either posterior or anterior segments of the medial temporal lobe (focused primarily on the hippocampus; see targeting in **Figure 2**).

In the task, context (the colour cue) was initially congruent with the touchscreen’s spatial location. Two touchscreens were attached to adjacent sides of the animal’s home-unit, separated by a corridor, and each assigned a different colour cue; yellow on the animal’s left, and blue on the animal’s right (**Figure 1A**). This congruence between colour cue and spatial location held during the Stable Context phase, but was later dissociated (see Unstable Context phase, below).

**Figure 1.**
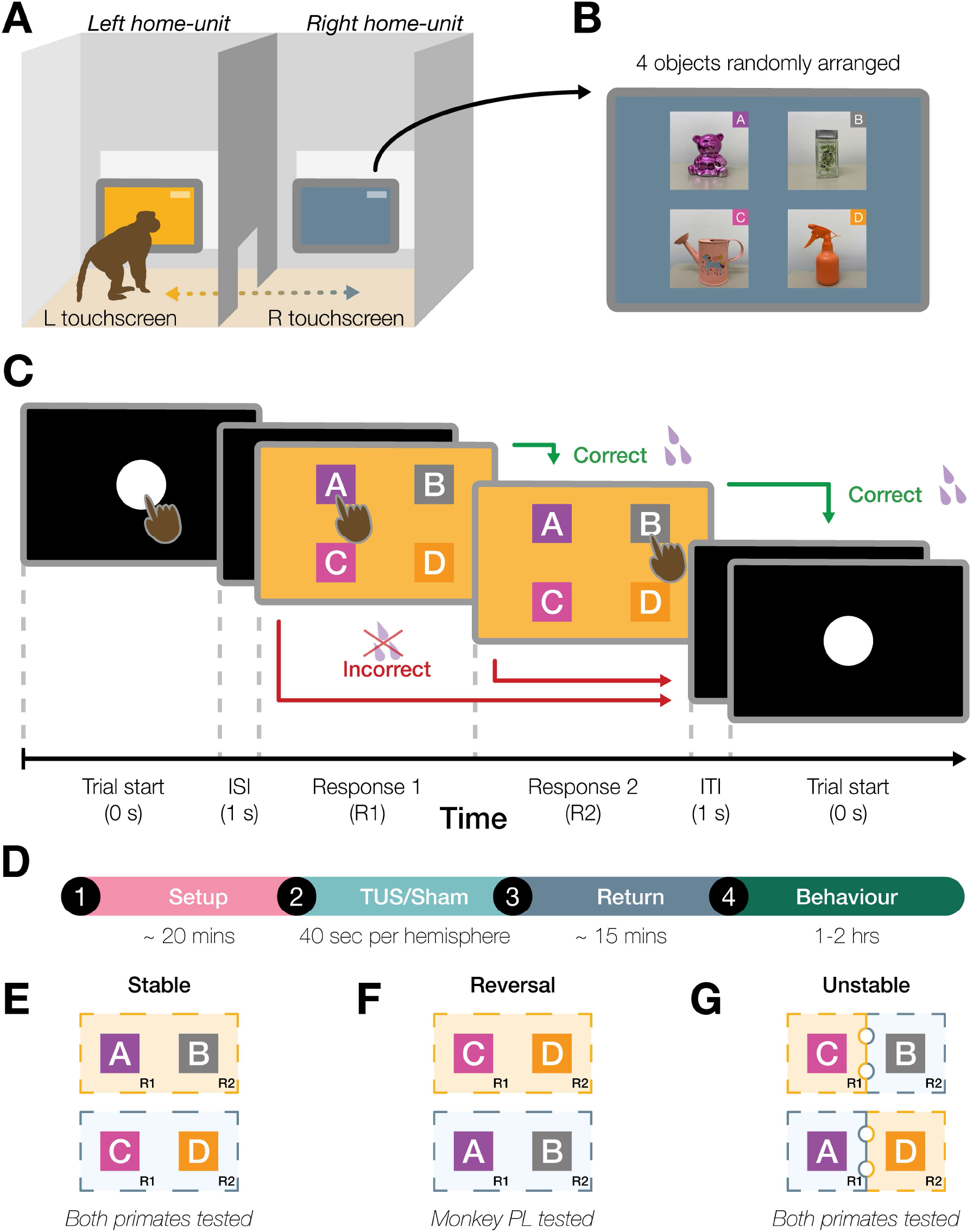
Schematic representation of the home-unit setup, trial sequence, and session timeline, along with the individual experimental phases. **A)** The touchscreen setup consisted of two adjacent rooms within the animal’s home-unit connected by a central corridor. Each home-unit contained a touchscreen within reach of the animal. **B)** Displays the four stimuli images used. Images were randomly arranged each trial. **C)** Depicts the time-course and sequence of events in a typical trial. Abbreviations: Inter-stimulus interval (ISI); Inter-trial Interval (ITI). **D)** Time course of a typical testing session, with animals being taken to the lab where TUS or sham stimulation to the medial prefrontal cortex, anterior and posterior medial temporal lobe (primarily the hippocampus; see Fig.2 and Supp. Fig.2) was applied, before returning the macaque to their home-unit to perform the behavioural task. **E)** Stable Context phase: the left touchscreen consistently displayed yellow-context trials (correct sequence: object A followed by object B), while the right touchscreen displayed only blue-context trials (correct sequence: object C followed by object D). **F)** Reversal Learning phase: involved a switch of the correct object sequences associated with each context. **G)** Unstable Context phase: the context changed mid-trial, and the macaque was expected to accommodate the change of context and temporal sequence (a correct response was the second item in the sequence associated with the context on the screen). During the Unstable Context phase, during each testing run, both stable and unstable context trials were randomly presented.

The active screen alternated every five trials, irrespective of performance, to ensure that the monkey had to regularly alternate between the two touchscreen locations. In a given trial, four object images (adapted from real-world objects used in a previous rodent task (Slater et al., 2025)) were randomly ordered onscreen (**Figure 1B**). Monkeys were trained via operant conditioning with juice reward to remember and to recall the correct context-specific two-object sequence. An example trial is illustrated in **Figure 1C**.

Prior to each home-unit testing session, TUS or sham (no-TUS stimulation) conditions were applied in the laboratory with the macaque in a testing chair to which it had been previously acclimatised. Each daily testing session began with TUS stimulation delivered to one of three prefrontal-hippocampal targets (**Figure 2**) or with a sham condition that simulated active stimulation at the same target sites but without delivering ultrasound. The TUS/sham and the three stimulation targets were counterbalanced across testing sessions. In the sham condition, the transducer was placed over the stimulation area, but the TUS transducer coil was not activated. Otherwise, the procedure was identical for both TUS and sham conditions. Stimulation (TUS or sham) was applied for 40 seconds per hemisphere while the macaque was awake, with the head temporarily immobilised for precise TUS targeting. Following TUS or sham, the animal was returned to its home-unit and was free to perform the touchscreen task for the next 1-2 hours without humans being present. **Figure 1D** outlines the timeline of an example testing session.

**Figure 2.**
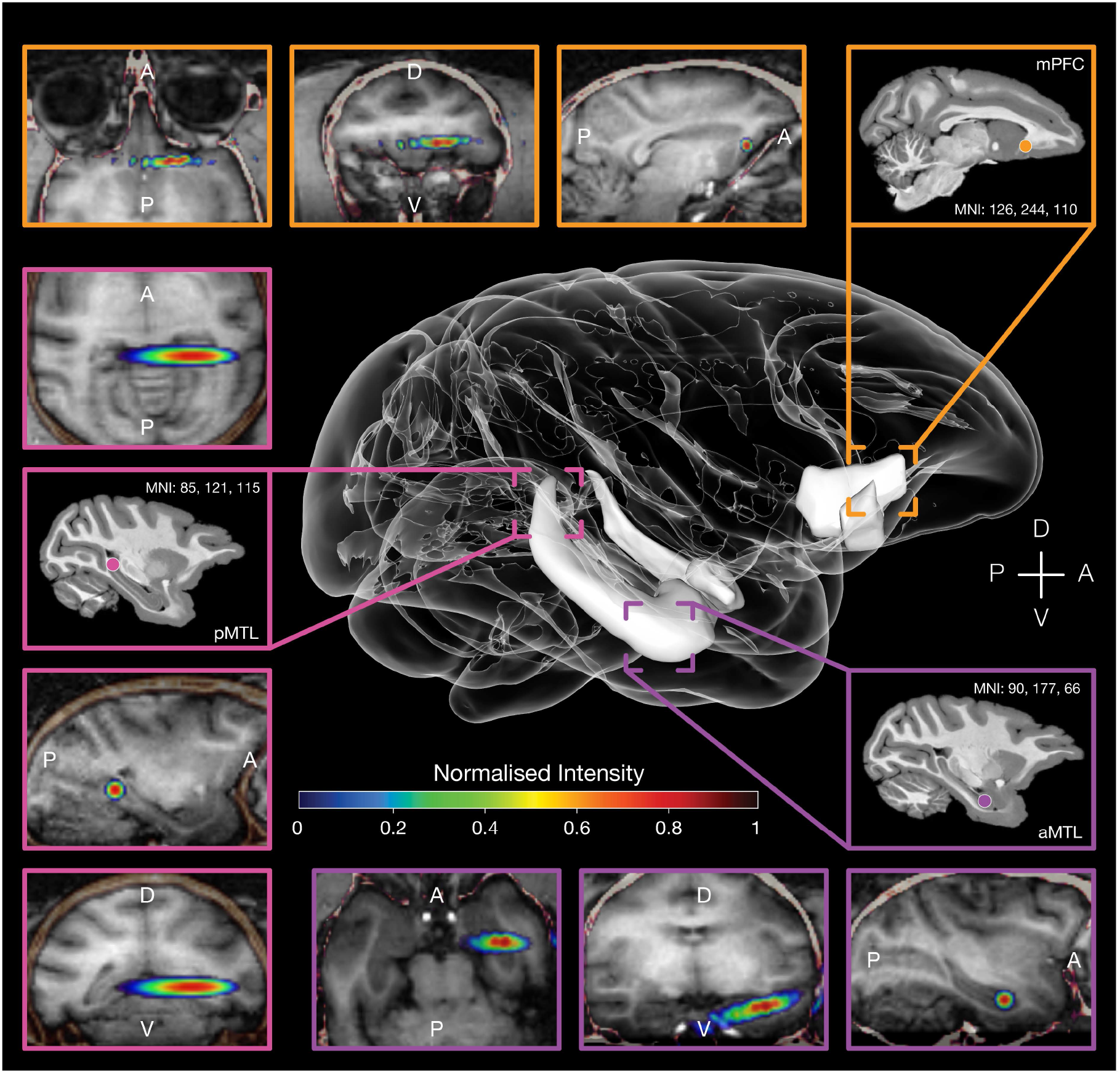
Ultrasound stimulation simulations. The figure illustrates the TUS simulations assisting the TUS targeting, conducted with k-plan software, modelled using Monkey PL’s MRI. Simulations for each of the three brain region targets: medial prefrontal cortex (outlined in orange); posterior medial temporal lobe (mostly the hippocampus; outlined in pink) and anterior medial temporal lobe outlined in purple. Individual coordinates in the macaque template MNI space correspond to each targeted area. Intensity of the sonication here has been normalised to highlight areas where intensity was at its peak.

Two rhesus macaques (1 male, Monkey PL; 1 female, Monkey MC) completed the full protocol. Unless otherwise noted, results reflect patterns statistically consistent and in the same direction of effect in both animals. Task performance was analysed using bias-reduced binomial logistic regression, with trial outcome (correct vs. incorrect) as the dependent variable and stimulation site (aMTL, pMTL, mPFC, and sham) as a categorical predictor. Sham trials served as the reference condition.

### Testing TUS impact on performance across stable contexts

In the initial, Stable Context phase, each context was consistently paired with a specific touchscreen (yellow on the left-side and blue on the right-side, **Figure 1E**). Monkeys were required to recall a fixed two-object sequence for each context (A → B for yellow, C → D for blue). The four objects were always presented in a random location on the screen.

Monkey PL completed sessions in a block design (several repeated stimulation sessions of one region with interleaved sham sessions). Monkey MC experienced fully counterbalanced stimulation and sham sessions across regions. For Monkey PL, the experimental blocking/ordering was an analytical factor.

Across both animals, stimulation of the anterior MTL reliably increased performance relative to sham. For Monkey PL, accuracy rose significantly from 91.7 % (± 2.09 SEM) during sham sessions to 94.1 % (± 1.59 SEM) following TUS directed at the aMTL (*ß* = 0.37 ± 0.11, *z* = 3.25, *p* = 0.001; **Figure 3A-B**). Monkey MC also showed a significant increase in performance from sham session performance of 89.1% (± 1.14 SEM) to 92.7% (± 1.11 SEM) with TUS of the anterior MTL (*ß* = 0.41 ± 0.20, *z* = 2.16, *p* = 0.031; **Figure 3C-D**). These convergent results suggest that anterior MTL TUS increased performance in both monkeys under the stable contextual environments.

**Figure 3.**
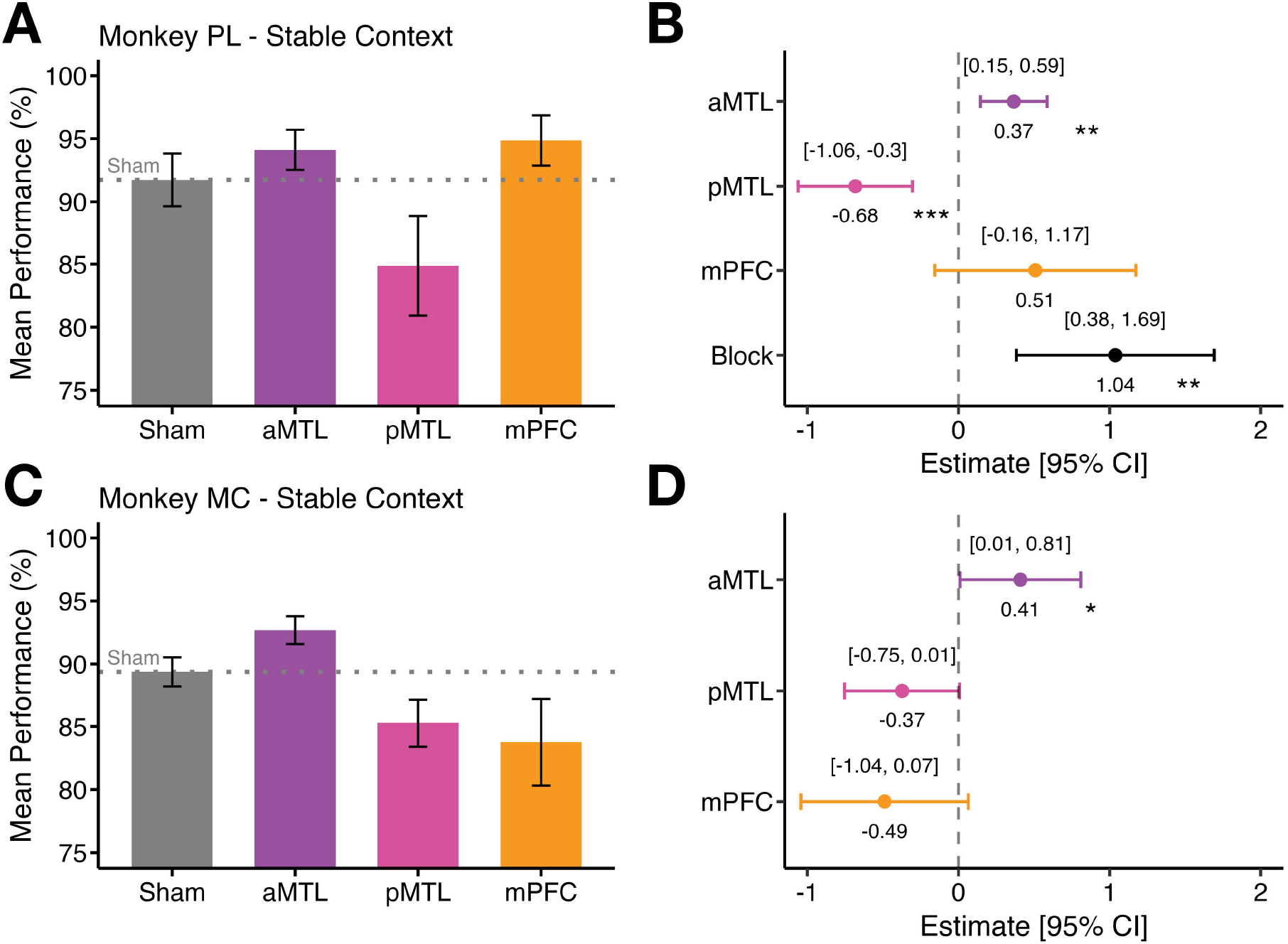
Effects of TUS of the three brain regions on task performance compared to sham across the Stable Context and Reversal Learning phases. **(A-D)** Stable Context phase: **A)** Mean performance (%) for sham, anterior medial temporal lobe (aMTL), posterior medial temporal lobe (pMTL), and medial prefrontal cortex (mPFC) stimulation in Monkey PL. Error bars represent the standard error of the mean (SEM). **B)** Logistic regression coefficient estimates with 95 % confidence intervals (CI) segregated by brain region in Monkey PL. The x-axis displays regression estimates, with the dashed line at 0 indicating no effect. **C)** Mean performance (%) with SEM bars for Monkey MC across the same brain regions. **D)** Logistic regression coefficient estimates with 95 % CIs segregated by brain region in Monkey MC.

By contrast, posterior MTL stimulation significantly reduced performance in one of the animals (PL) and was a statistical trend in the same direction for the other (MC). Namely, reduced performance with pMTL TUS was significant in Monkey PL (pMTL TUS: 84.9 % ± 3.95 SEM vs. sham: 94.1 % ± 1.59 SEM; *ß* = - 0.68 ± 0.19, *z* = - 3.53, *p* < 0.001; **Figure 3A-B**), and showed a statistical trend in the same direction in Monkey MC (pMTL TUS: 85.2 % ± 1.84 SEM vs. sham: 92.7% ± 1.11 SEM; *ß* = - 0.37 ± 0.19, *z* = - 1.90, *p* = 0.057; **Figure 3C-D**).

Stimulation of the medial prefrontal cortex during the stable context task performance did not significantly alter accuracy in either monkey. For Monkey PL, performance following TUS to the mPFC did not significantly differ (94.9 % ± 1.99 SEM) compared to sham (94.1 % ± 1.59 SEM; *ß* = 0.51 ± 0.34, *z* = 1.50, *p* = 0.134; **Figure 3A-B**). And in Monkey MC, performance following mPFC stimulation (83.8 % ± 3.47 SEM) showed only a statistical trend in reduced performance compared to sham performance (92.7% ± 1.11 SEM; *ß* = - 0.49 ± 0.28, *z* = - 1.65, *p* = 0.098; **Figure 3C-D**).

In Monkey PL, the block design revealed a main effect of session block (*ß* = 1.04 ± 0.33, *z* = 3.11, *p* = 0.002; **Figure 3A-B**), with performance improving over the course of testing, importantly regardless of stimulation condition: TUS or sham. For Monkey MC, there was no evidence of a systematic effect of session date and including it as either a fixed or random effect did not improve model fit (Δ AIC < 2). Accordingly, we report the simpler model with only stimulation region as a predictor (**Figure 3C-D**).

Together, these findings reveal a functional dissociation that was consistent in both animals: TUS of the anterior MTL led to improved sequence recall performance when contexts were stable, which was not the case for TUS of the posterior MTL (opposite effect – significant in one monkey and a statistical trend in the other) or the mPFC (no significant effect in one monkey; opposite effect statistical trend in the other).

### Testing TUS impact on performance during unstable contexts

We next tested the impact of TUS on task performance under unstable contextual conditions. In the previous phases, sequence memory was tested in stable contexts, in which the background colour presented on the touchscreen remained constant throughout the trial and was consistently associated with a specific object sequence and touchscreen. In contrast, during the Unstable Context phase, contextual uncertainty was increased by systematically introducing a context switch mid-trial: following the selection of the first correct item, the background colour could change to indicate a different context before the second choice was required. Critically, this manipulation required the macaque to update the contextual information while maintaining the temporal position within the ongoing sequence. Under these conditions, the correct choice was the item occupying the second position in the sequence appropriate for the updated context. Importantly, the total reward magnitude and number of rewarded responses per trial were identical to those in the stable context sessions; only the contextual decision rule was altered. Moreover, any context could occur on either of the two touchscreens, removing the spatial location of the touchscreen as a reliable contextual cue (**Figure 1G**).

Across both animals, mPFC stimulation significantly increased performance relative to the sham conditions. Monkey PL’s accuracy increased from 77.5% during the sham condition (± 0.79 SEM) to 85.5 % ± 0.86 SEM with active TUS to mPFC (*ß* = 0.54 ± 0.08, *z* = 6.47, *p* < 0.001; **Figure 4A-B**). Similarly, Monkey MC showed significantly improved performance with mPFC TUS (80.5 % ± 1.46 SEM) relative to sham (73.5 % ± 1.37 SEM; *ß* = 0.40 ± 0.12, *z* = 3.41, *p* < 0.001; **Figure 4C-D**).

**Figure 4.**
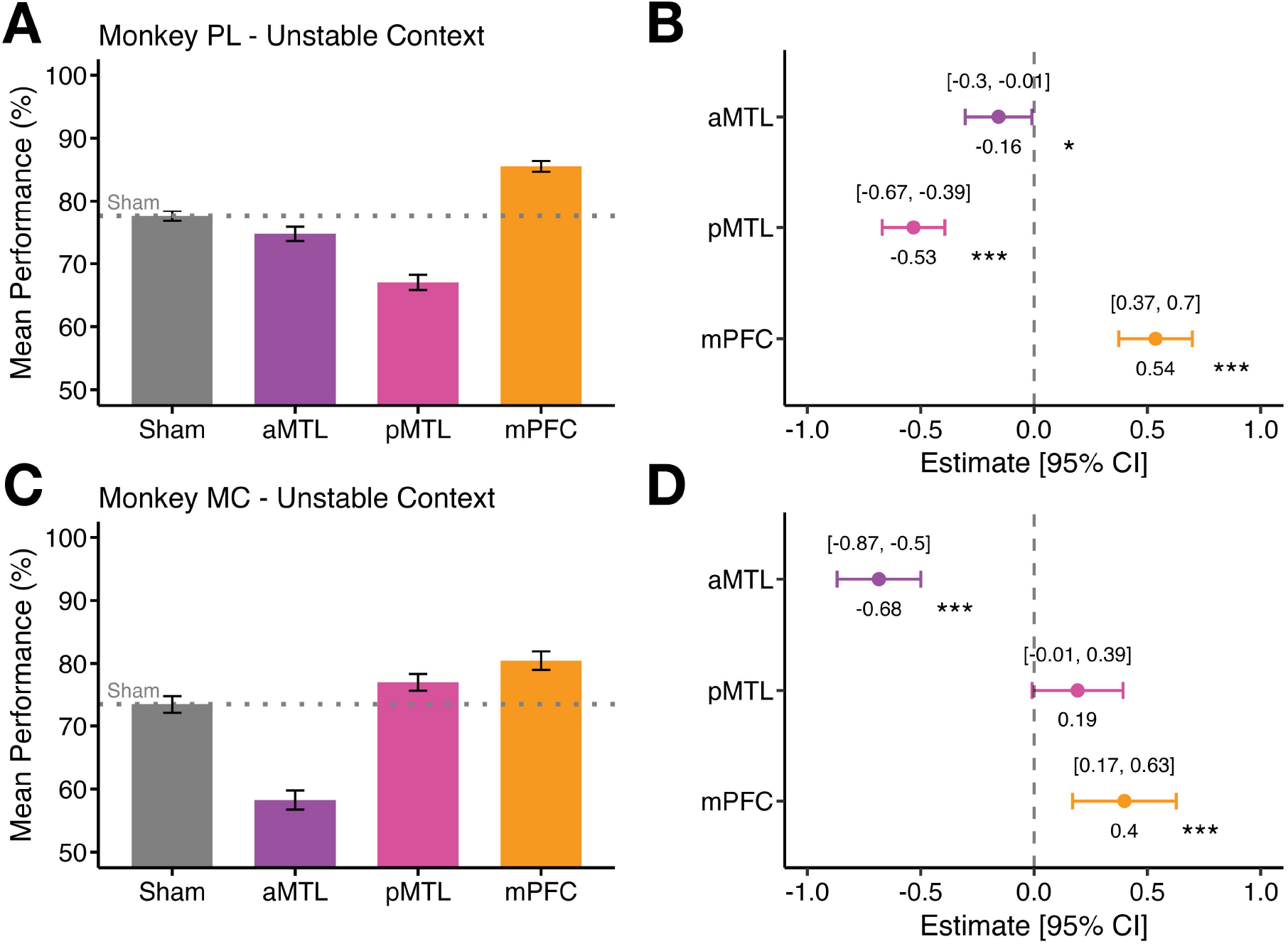
Effects of stimulation on trial performance in three brain regions compared to sham for the Unstable Context phase. **A)** Mean performance (%) with standard error of the mean (SEM) bars for sham, aMTL, pMTL, and mPFC stimulation relative to sham in Monkey PL. Error bars represent the standard error of the mean (SEM). **B)** Logistic regression coefficient estimates with 95 % confidence intervals (CI) segregated by brain region in Monkey PL. The x-axis displays regression estimates, with the dashed line at 0 indicating no effect (sham condition levels). **C)** Mean performance (%) with SEM bars for Monkey MC across the same brain regions. **D)** Logistic regression coefficient estimates with 95 % CIs segregated by brain region in Monkey MC.

By contrast, aMTL stimulation significantly impaired performance in both animals relative to sham. Monkey PL’s performance significantly dropped from 77.5% ± 0.79 SEM during sham conditions to 74.8 % ± 1.13 SEM following aMTL TUS (*ß* = - 0.16 ± 0.08, *z* = - 2.09, *p* = 0.037; **Figure 4A-B**). Monkey MC’s performance also significantly worsened following aMTL stimulation from 73.5 % ± 1.37 SEM during sham conditions to 58.3 % ± 1.52 SEM (*ß* = - 0.68 ± 0.09, *z* = - 7.26, *p* < 0.001, **Figure 4C-D**).

Posterior MTL TUS showed inconsistent opposing results in the two monkeys with Monkey PL demonstrating a significant reduction in accuracy (67.0 % ± 1.20 SEM) compared to sham (77.5% ± 0.79 SEM; *ß* = - 0.53 ± 0.07, *z* = - 7.54, *p* < 0.001; **Figure 4A-B**), but Monkey MC displaying a positive statistical trend following pMTL stimulation (77.0 % ± 1.31 SEM) compared to sham (73.5 % ± 1.37 SEM; *ß* = 0.19 ± 0.10, *z* = 1.88, *p* = 0.061; **Figure 4C-D**).

In summary, although aMTL TUS improved performance under the Stable Context conditions (see **Figure 3** above), mPFC TUS improved performance under the Unstable Context. Under either Stable or Unstable Contexts, TUS of the aMTL or pMTL, respectively, either disrupted or did not have a consistent effect on performance in both macaques.

### Testing TUS impact on performance during sequence relearning (Monkey PL only)

In Monkey PL, we were able to test the impact of site-specific TUS on performance during *Reversal Learning*. In the *Reversal Learning* phase, the two-object sequences associated with each context were reversed. Specifically, objects A and B, previously linked with the yellow context, were assigned to blue, and objects C and D assigned to yellow (**Figure 1F**). Sessions were counterbalanced with sham stimulation, allowing direct comparison across brain regions.

Stimulation of the aMTL significantly improved performance relative to sham (active TUS: 89.8 % ± 0.75 SEM vs sham: 78.3 % ± 0.80 SEM; *ß* = 0.89 ± 0.09, *z* = 9.42, *p* < 0.001; **Figure 5A-B**). Medial prefrontal cortex stimulation also improved accuracy (TUS: 85.0 % ± 0.85 SEM vs sham: 78.3 % ± 0.80 SEM; *ß* = 0.451 ± 0.082, *z* = 5.53, *p* < 0.001, **Figure 5A-B**). As reported previously under the stable context above, TUS to the pMTL did not significantly alter performance (TUS: 76.8 % ± 1.07 SEM vs sham: 78.3 % ± 0.80 SEM; *ß* = - 0.09 ± 0.08, *z* = -1.15, *p* = 0.249, **Figure 5A-B**). The same test in Monkey MC was not possible before the animal completed all studies; therefore, this individual animal result is reported only within the context of monkey PL’s individual performance.

**Figure 5.**
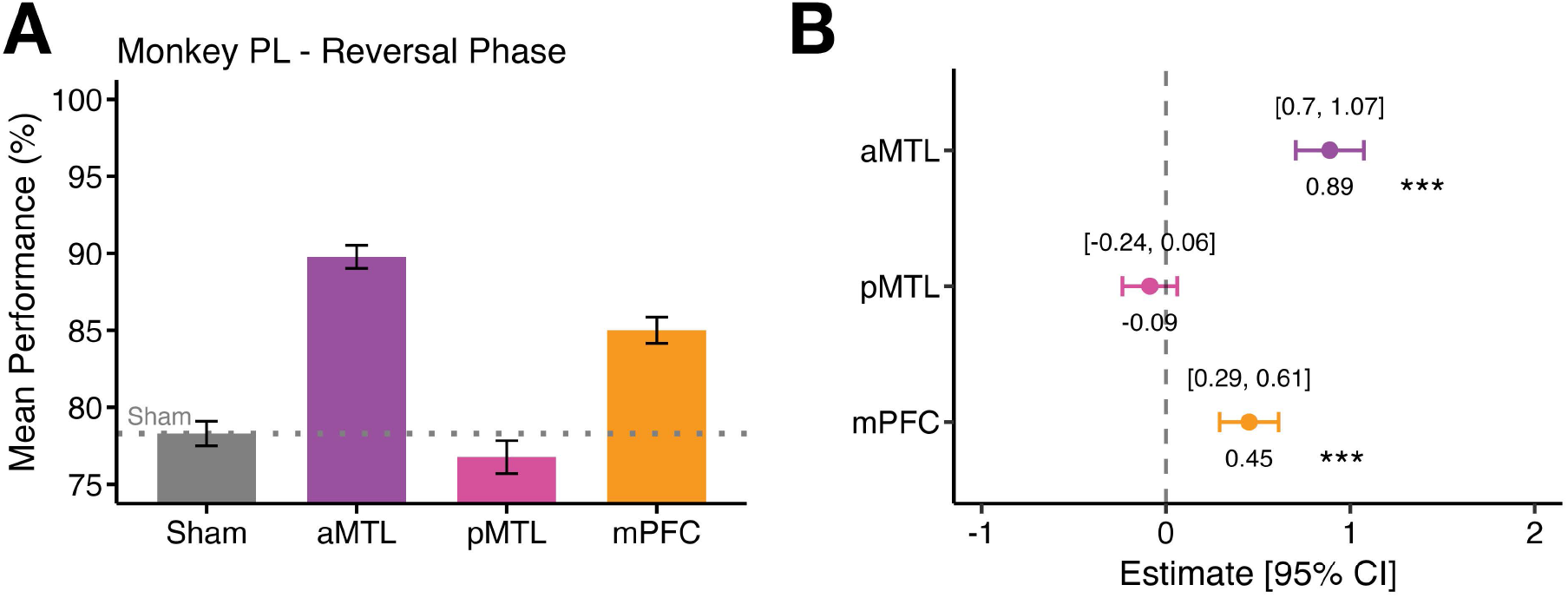
Effects of stimulation on trial performance in three brain regions compared to sham for the Reversal phase (tested only in Monkey PL). **A)** Mean performance (%) with standard error of the mean (SEM) bars for sham, aMTL, pMTL, and mPFC stimulation relative to sham in Monkey PL. Error bars represent the standard error of the mean (SEM). **B)** Logistic regression coefficient estimates with 95 % CIs segregated by brain region in Monkey PL. Significance is indicated by * (p < .05), ** (p < .01) or *** (p < .001) following binomial logistic regression.

### Insights from between monkey testing variability

Two key methodological differences in how the monkeys were tested provide important insights for interpreting the results. For the first primate tested (Monkey PL), we initially adopted a blocked design of TUS and sham sessions, as it was unclear whether measurable effects would require grouping across sessions. Post-hoc analyses revealed a main effect of session block (*ß* = 1.04 ± 0.33, *z* = 3.11, *p* = 0.002; **Figure 3A-B**), with performance improving over the course of testing. Crucially, however, including session block in the model did not alter the effects of stimulation regions. To avoid potential temporal confounds, we switched to a fully counterbalanced session-by-session design for Monkey MC and for the remainder of Monkey PL’s testing (*Reversal* and *Unstable Context* phases), ensuring that any learning-related improvements could be more directly aligned with sham sessions conducted on adjacent days. For Monkey MC, modelling session date as either a fixed or random effect revealed no change in performance over time (Δ AIC < 2), and we therefore report the simpler model including only stimulation region as a predictor (**Figure 3C-D**).

The second facet in the performance data was that the two monkeys had different preferences in how they were tested that required constraining the testing approach. Monkey MC was more reluctant to move between the two testing rooms of their home-unit and risked providing little data to evaluate behavioural performance under the different TUS and sham conditions. By comparison, Monkey PL continued to alternate between the familiar two home-units every five trials (**Figure 6A**), whereas Monkey MC would only perform the testing sessions on a single touchscreen, with side of home-unit testing counterbalanced across sessions (**Figure 6B**). For Monkey MC, this included the familiar yellow home-unit and an unfamiliar home-unit, that had not been associated with any context or trial type during training or testing.

**Figure 6.**
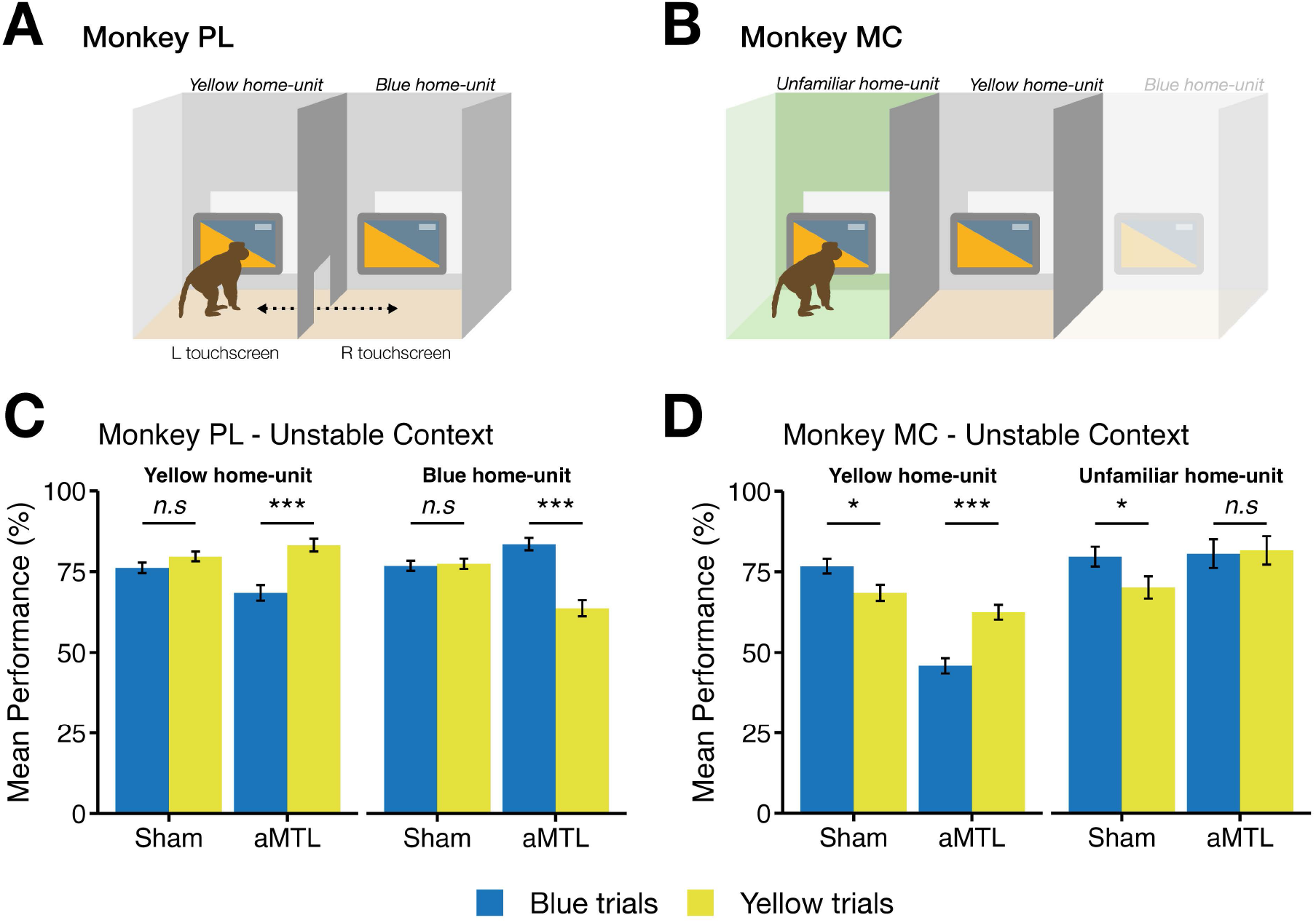
Effects of home-unit and context on accuracy during the Unstable Context phase. **A)** Task setup for monkey PL: sessions involved alternating between the yellow and a blue home-unit every 5 trials, with each home-unit previously associated with its respective context. **B)** Task setup for monkey MC: each session was restricted to a single home-unit, either the familiar yellow home-unit or a novel unfamiliar home-unit not encountered during training. **C)** Mean performance (± SE) for monkey PL for yellow- and blue-context trials when tested in either the yellow or blue home-unit. **D)** Mean performance (± SE) for monkey MC on yellow- and blue-context trials when tested in either the yellow or the unfamiliar home-unit. Data is shown separately for sham sessions (no TUS) and aMTL sessions (anterior medial temporal lobe TUS). Significance is indicated by *** (p < .001), * (p < .05) or n.s (p > .05).

To examine how these differences influenced performance, we analysed trials as a function of home-unit side and context, revealing an intriguing home-unit-dependent effect of aMTL stimulation relative to sham, as follows.

For Monkey PL, aMTL TUS enhanced performance specifically when the trial’s context matched the home-unit previously associated with that context. Following aMTL stimulation, accuracy was high for yellow-context trials in the yellow home-unit (83.2 % ± 1.95 SEM) and blue-context trials in the blue home-unit (83.5 % ± 1.92 SEM). In contrast, performance dropped when the trial context did not match the home-unit association: blue-context trials in the yellow home-unit (68.4 % ± 2.43 SEM) and yellow-context trials in the blue home-unit (63.7 % ± 2.48 SEM). Pairwise comparisons confirmed these interactions: in the yellow home-unit, yellow-context trials were significantly more accurate than blue-context (*ß* = 0.83 ± 0.18, *z* = 4.59, *p* < 0.001; **Figure 6C**), and in the blue home-unit, blue-context trials were significantly more accurate than yellow-context trials (*ß* = 1.06 ± 0.18, *z* = 6.02, *p* < 0.001; **Figure 6C**).

Sham stimulation produced no such bias for either side. Yellow-context trials (79.7 % ± 1.50 SEM) were performed similarly to blue-context trials (76.2 % ± 1.63 SEM) in the yellow home-unit (*ß* = - 0.21 ± 0.13, *z* = - 1.59, *p* = 0.111; **Figure 6C**). Likewise, blue-context trials (76.8 % ± 1.57 SEM) were performed at similar accuracy level to yellow-context trials (77.4 % ± 1.60 SEM), even when experienced in the blue home-unit during a sham session (*ß* = - 0.04 ± 0.13, *z* = - 0.29, *p* = 0.77; **Figure 6C**).

Monkey MC showed a comparable pattern with aMTL stimulation significantly increasing performance for yellow-context trials (62.5 % ± 2.29 SEM) compared to blue-context trials (45.9 % ± 1.60 SEM) when performed in the yellow home-unit (*ß* = 0.68 ± 0.14, *z* = 4.96, *p* < 0.001; **Figure 6D**). *Sham* performance, however, showed an opposite effect: significantly increased performance for blue-context trials (76.8 % ± 2.29 SEM) compared to yellow-context trials (68.5 % ± 2.51 SEM) when in the yellow home-unit (*ß* = 0.42 ± 0.17, *z* = 2.43, *p* = 0.015; **Figure 6D**).

This effect of context-specific improvement in performance related to Monkey MC’s preference for the blue/yellow context persisted for Monkey MC when they were in the unfamiliar home-unit, with blue-context trials (79.7 % ± 3.05 SEM) being significantly more accurate than yellow-context trials (70.2 % ± 3.46 SEM; *ß* = 0.51 ± 0.25, *z* = 2.05, *p* = 0.041; **Figure 6D**). What is striking, however, is that the aMTL effect, previously seen in Monkey PL and Monkey MC, disappeared when Monkey MC performed trials in the unfamiliar home-unit, with both yellow-context (81.6 % ± 4.38 SEM) and blue-context trials (80.6 % ± 4.44 SEM) being performed equally well (*ß* = 0.07 ± 0.41, *z* = 0.16, *p* = 0.870; **Figure 6D**). These observations indicate that when context-home-unit associations were maintained, aMTL stimulation boosted accuracy, whereas in unfamiliar environments the effect of aMTL TUS on stable context performance was absent, providing further insights into the animals’ preferences, performance and site-specific TUS effects on performance.

## DISCUSSION

The present study provides initial evidence for selective enhancement of cognitive function using low-intensity, offline Transcranial Ultrasound Stimulation (TUS) conducted with two nonhuman primates performing a novel context-guided memory sequencing task across multiple touch screens in their home units. Although the sample size is necessarily small (two monkeys with different training histories and testing preferences – see Results), only statistically consistent effects are reported in both animals following stimulation of specific medial temporal lobe (MTL) and prefrontal targets. Stimulation of the anterior MTL (aMTL), focused on and largely affecting the anterior hippocampus (aHPC), see **Supplementary Figure 2**, enhanced performance under stable contextual conditions. By contrast, stimulation of the medial prefrontal cortex (mPFC) improved performance under unstable conditions that required the macaques to adapt to contextual changes, and aHPC stimulation disrupted performance under the unstable context conditions.

Performance improvement or disruption was not observed in the macaques with stimulation of posterior MTL (pMTL), centred on the posterior hippocampus (pHPC), or with sham stimulation. These results provide the first evidence that low-intensity TUS can selectively bias cognitive function and, in so doing, identify the positive causal elements of adaptable cognition in fronto-temporal systems.

### aMTL Stimulation Enhanced Memory Sequencing in Stable Contexts

The differential behavioural effects of anterior versus posterior medial temporal lobe stimulation likely reflect the distinct functional specialisations along the hippocampal longitudinal axis contributing to memory functions. Across species, the anterior hippocampus has been associated with the retrieval of temporal sequences (Lehn et al., 2009) and with the contextualised representations, associating episodic memories within the environmental scenario within which they occur (Sheldon and Levine, 2016).

By comparison, the posterior hippocampus is associated with the retrieval of local spatial and perceptual mnemonic content (Poppenk et al., 2013; Zeidman and Maguire, 2016). Under conditions where the colour cue (context) remained stable and congruent with spatial location, both reliably predicting the correct sequence, aMTL stimulation improved task performance in both animals. Entering a particular home-unit (e.g., the left unit consistently associated with yellow-context trials) in combination with TUS to the aMTL appeared to facilitate retrieval of the corresponding memory sequence in both macaques, consistent with evidence that re-entering a familiar context primes relevant memories and facilitates retrieval (Smith and Bulkin, 2014). In contrast, stimulation of the pMTL did not produce reliable improvement or impairment. However, when the context changed mid-trial and spatial cues no longer aligned with the context, aMTL stimulation impaired performance in both monkeys, suggesting that the same facilitative process that benefits stable contextual retrieval may interfere with the recall of previously learned associations under a different context when the environmental cues become unreliable.

These observations provide bidirectional causal perturbation evidence in support of the interpretation that aMTL stimulation can strengthen the reliance on contextual representations, either supporting or hindering performance depending on the stability of the environmental context guiding memory retrieval.

### pMTL Stimulation Did Not Affect Performance

Stimulation of the pMTL did not reliably influence performance under either stable or unstable contexts, although given the focality of stimulation we cannot rule out that a more effective pMTL locale was not stimulated. Although posterior hippocampal involvement in fine-grained spatial memory is well supported in other species (Hoscheidt et al., 2010; Nadel et al., 2013), the present task may have not sufficiently engaged the processes most reliant on the pMTL site that was stimulated. We did observe significantly reduced performance under the stable context during pMTL stimulation in one animal, which was a statistical trend in the same direction in the other animal, but we avoid interpreting this result out of an interest to only interpret effects that were significant and consistent in both animals.

### mPFC Stimulation Facilitated Adapting to Contextual Change

In contrast to the aMTL stimulation effects described above, which resulted in significantly impaired performance under the unstable contexts in both animals, stimulation of the mPFC selectively enhanced performance under the unstable context conditions in the animals. This provides positive causal perturbation evidence consistent with the proposed role of the prefrontal cortex in resolving interference between overlapping memories and guiding hippocampal retrieval under changing environmental conditions (Euston et al., 2012; Navawongse and Eichenbaum, 2013; Preston and Eichenbaum, 2013). When contextual contingencies change, we speculate that mPFC stimulation may have aided disambiguation of competing sequence representations, assisting in coordinating hippocampal retrieval processes under the new context (Milivojevic et al., 2015). This interpretation is consistent with the involvement of mPFC in schema updating and adaptive mechanisms that accommodate changes in environmental contexts during memory recall (Kesteren et al., 2012; McKenzie et al., 2014; Tse et al., 2007).

### Interpreting Enhancement Effects Within the Broader Network

Of our two alternative hypotheses, the one that stipulated that low-intensity TUS would enhance rather than disrupt memory performance was supported and is the more parsimonious interpretation of the results. The less parsimonious explanation is that stimulation of the aMTL or mPFC locally disrupted the function of these nodes. In this case, enhancement of performance under the, respectively, stable and unstable context would depend on compensation by the other nodes in the system. However, this poses a more complex system compensatory explanation that would be at odds with the known roles of the aMTL, pMTL and PFC in context-guided memory. Nonetheless, network-level effects are expected, and resting-state functional MRI data collected from macaques with a similar offline TUS protocol shows how TUS modulates both local and distributed network connectivity (Folloni et al., 2019; Verhagen et al., 2019). Moreover, we recently reported on the longevity of left aMTL stimulation in Monkey PL under this study, which showed the tens of minutes long time course of TUS effects on the broader system. That study showed increases in functional connectivity and increases in local network interactions in prefrontal cortex lasting for over an hour after stimulation, with decreases in spontaneous network fluctuations, suggesting that hippocampal stimulation can initiate a reorganisation of large-scale functional networks (Atkinson-Clement et al., 2025). Although the entorhinal cortex (ERC) was not as strongly stimulated as the hippocampus in our study (**Supplementary Figure 2**), electrical stimulation of the ERC has been shown to rescue spatial memory in mouse models of Alzheimer’s disease (Mann et al., 2018) and enhances spatial memory during learning in human epilepsy patients (Suthana et al., 2012).

### Comparison to Other Brain Stimulation Studies

The intensities used in this study fall within the low-intensity, non-thermal range validated in nonhuman primate work (Bongioanni et al., 2021; Khalighinejad et al., 2020; Verhagen et al., 2019), distinguishing the present approach from direct moderate-intensity, microbubble-assisted ultrasound used in Alzheimer’s model rodents (Blackmore et al., 2023). For instance, blood-brain barrier opening with TUS has been shown to elicit neurotrophic effects and memory enhancement in rodents performing in the Morris water maze (Shen et al., 2020), via upregulation of brain-derived neurotrophic factor (BDNF) a mediator of neurogenesis and synaptic plasticity (Shin et al., 2019). At the cellular level, TUS-generated mechanical pressure waves modulate mechanosensitive ion channel gating, capable of increasing neuronal excitability, although longer lasting effects on metabotropic ion channels, such as NMDA, may be indirect via interaction of neurons with astrocytes (Kubanek and Jan, 2018; Yoo et al., 2022). In hippocampal slice cultures, direct low-intensity TUS effects can enhance CA1 pyramidal neuron activity (Tyler et al., 2008), and repeated exposure (over 10 days) increases dendritic spine density and glutamate receptor expression (GluN2A subunits), enhancing spontaneous excitatory post-synaptic currents (Huang et al., 2019). Moreover, TUS in rodents combined with two photon calcium imaging has identified specific excitatory and inhibitory neuronal activation effects (Lemaire et al., 2024).

The offline TUS stimulation parameters used here (intensity: 10 W/cm^2^; duty cycle: 30 %; 10 Hz pulse frequency) fall within the range predicted by a computational model of TUS effects to favour enhancement rather than suppression of neuronal activity, although as we have noted that model remains controversial and current predictability in the directionality of TUS effects remains low (Caffaratti et al., 2025). Computational modelling notwithstanding, rodent intracranial recordings indicate that our combination of pulse-repetition frequency, duty cycle, and stimulation intensity should generally produce excitatory effects on neurons in the stimulated brain, at least for cortical areas (Kim et al., 2024; Murphy et al., 2024).

The 10 Hz pulse-repetition frequency used here falls within the alpha band (8-13 Hz) rather than the theta band (4-8 Hz). Interestingly, the lower edge of the 10-15 Hz alpha range has recently been shown to increase in magnitude in hippocampal-connected cortex during retrieval of learned item-in-context associations in macaques (Hussin et al., 2022), in contrast to theta band (4-8 Hz) activity, which is commonly suppressed during item-in-context memory retrieval in these species (Abbaspoor et al., 2023). As such, the present protocol is more consistent with retrieval-related alpha-band dynamics than with the classically defined theta-band prefrontal-hippocampal synchrony.

How do we reconcile these findings with prior TUS studies reporting disruptive effects in nonhuman primates? Notably, disruptive effects have been reported using a nearly identical stimulation protocol to the one used here, targeting medial prefrontal cortex during value-based decision making, as reported by Fouragnan *et al*. (2019), Boroujeni *et al*. (2022), and Bongioanni *et al*. (2021). These prior results targeted the dorsal ACC, a region implicated in reward-based valuation and reinforcement learning.

By comparison, our mPFC target was Brodmann area 25, selected for its dense direct hippocampal input: area 25 receives the densest hippocampal projections of any prefrontal region (Barbas and Blatt, 1995), and is anatomically and functionally distinct from dorsal ACC. As such, the putatively discordant direction of effect across these nonhuman primate TUS studies might reflect a distinction in the brain networks and circuits being targeted, rather than a contradiction in the directionality of effects of TUS itself. Direct neural recordings during the task in nonhuman primates or neurosurgery patients, with and without TUS, will be required to address whether there is indeed a discrepancy or provide a better understanding on the basis of the variability in directionality of TUS effects. Nonetheless, our results raise the possibility that this TUS protocol can have enhancing behavioural effects, in concord with the rodent literature on TUS-induced neuronal excitation (Murphy and Fouragnan, 2024).

Most prior TUS studies in humans reporting behavioural effects have focused on direct (‘online’) TUS stimulation of sensory, motor, or higher-level visual areas (Butler et al., 2022; Fomenko et al., 2020; Legon et al., 2014; Liu et al., 2021). Few TUS studies have reported effects on cognitive function, and none, to our knowledge, have demonstrated the site-specific and selective enhancement of cognitive function shown here in a primate model. In a small study of eight Alzheimer’s patients, right hippocampal TUS was reported to have improved recall and recognition memory amidst a battery of neuropsychological assessments, although the absence of sham or control-site stimulation and the lack of correction for multiple comparisons limit interpretability (Jeong et al., 2022). More controlled work by Fine and colleagues has shown that online TUS to the right inferior frontal gyrus during the go cue presentation in a stop-signal task can improve stopping performance, which was not observed with somatosensory cortex stimulation in control participants (Fine et al., 2023), see also (Zhang et al., 2021). A human study applying TUS to dorsal ACC or anterior insula reported alterations in learned biases, including perseveration, but the authors note that they could not draw conclusions on whether the effects were excitatory or inhibitory, only that they were consistent in both brain areas (Koutsoumpari et al., 2026). We have also recently reported that theta-burst TUS to the anterior temporal lobe increases semantic memory performance in healthy participants, coinciding with reductions in GABA and increases in glutamate/glutamine signal levels in the target region measured with magnetic resonance spectroscopy (Jung et al., 2025). Importantly, emerging evidence indicates that ‘offline’ TUS can also have sustained cognitive effects: brief theta-frequency stimulation (80 s) has been shown to improve performance on demanding conditions of the stop-signal task for over an hour after stimulation (Atkinson-Clement et al., 2024), demonstrating that short-duration, low-intensity ultrasound can induce longer-lasting neuromodulatory changes.

### Limitations

The key limitation inherent to neuroscientific studies with nonhuman primates is the small sample size, which is required to ensure that the smallest possible number of nonhuman primates is used per the Reduction principle of the 3Rs (Petkov et al., 2022). Although the primates had different learning histories, which formed the basis of our further analysis of their behavioural strategies (see Results), we only report and interpret effects that were significant and consistent in directionality of effects by brain area in both animals. Also, while TUS is known for its precision in targeting deep brain structures, the cigar-shaped pulse cannot be focused specifically on any one area, and this pulse can stimulate both grey and white matter. Therefore, we do not attribute effects to any specific brain area within the anterior/posterior medial temporal lobe and medial prefrontal cortex. Within the MTL, the hippocampus was the most consistently affected brain area, and for the PFC, the target was Brodmann area 25, an area that is highly interconnected with the MTL. Although we cannot rule out stimulation of fibres of passage, evidence that TUS can modulate white matter is limited with a recent report showing ultrasound-induced plasticity in the human corticospinal tract using a 5 Hz stimulation protocol (Curtin et al., 2025).

### Summary

Taken together, these findings highlight dissociable contributions of the anterior medial temporal lobe and medial prefrontal cortex in context-dependent memory for sequences. Transcranial ultrasound to the anterior medial temporal lobe including hippocampus supports retrieval of global contextual associations, and the medial prefrontal cortex can help the cognitive system adapt to changes in environmental scenarios. Low-intensity transcranial ultrasound stimulation provides a non-invasive means to transiently bias these distinct functions, offering new positive causal impact insights into the mechanistic architecture of the fronto-temporal memory network.

## RESOURCE AVAILABILITY

Data reported in this paper will be made available on the Open Science Framework under https://osf.io/arqp8/ in the folder associated with this paper (Slater *et al*.).

## ACKNOWLEDGMENTS

This work was supported by MRC: MR/X003701/1, UKRI 527, EPSRC: EP/W004488/1, EP/X01925X/1, EP/W035057/1, and BBSRC: BB/T008695/1. CIP was supported by: Carver Trust Professorship, NIH (U01NS137991 & MH109429) & NSF (SBE2342847).

## SUPPLEMENTARY INFORMATION

**Supplementary Figure 1.**
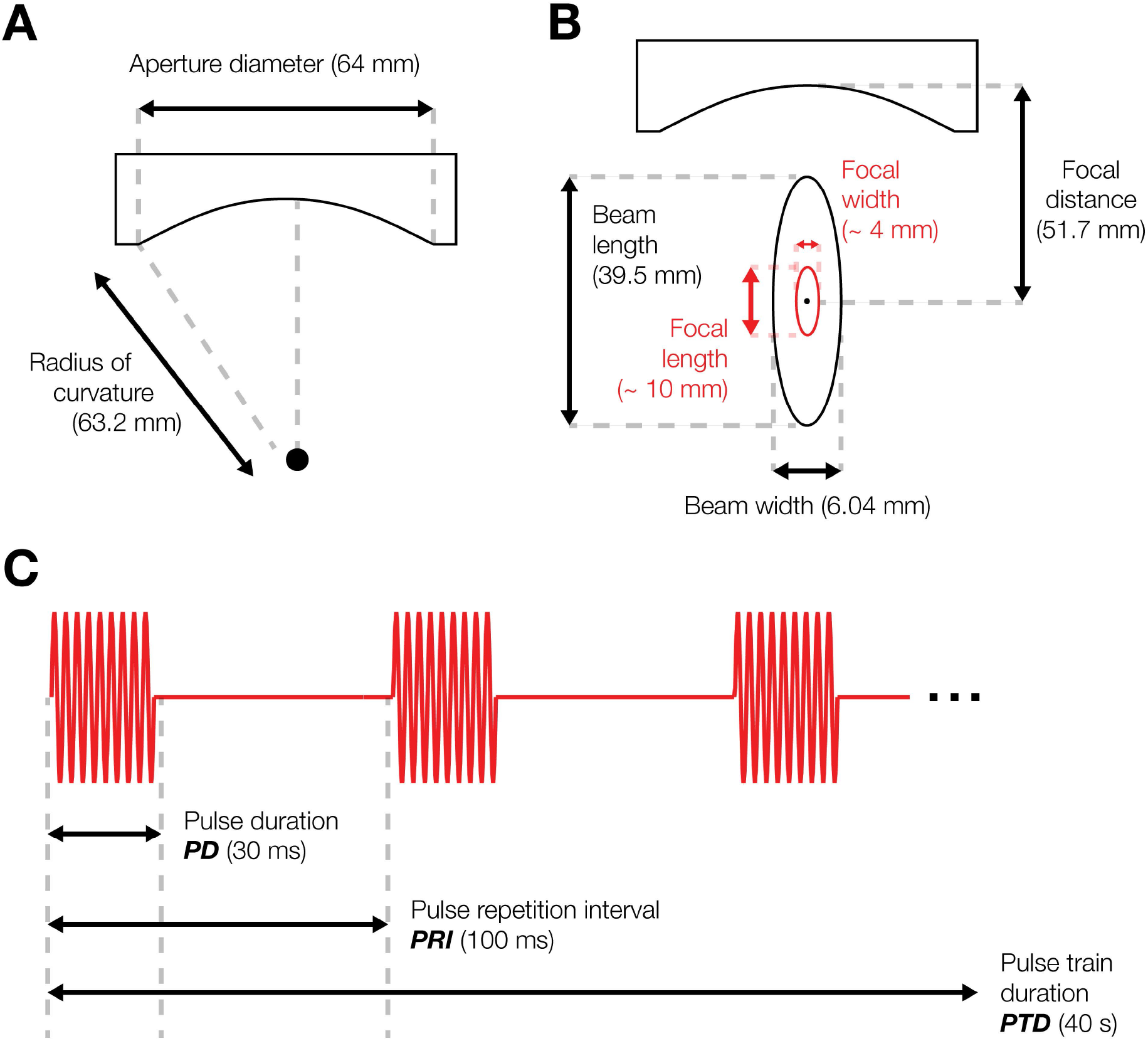
Transducer dimensions and waveform parameters as outlined by ITRUSST. **A)** Diameter of the transducer, and **B)** parameters of the focal distance, beam length, and beam width. **C)** Schematic of the ultrasound pressure waveform.

**Supplementary Figure 2.**
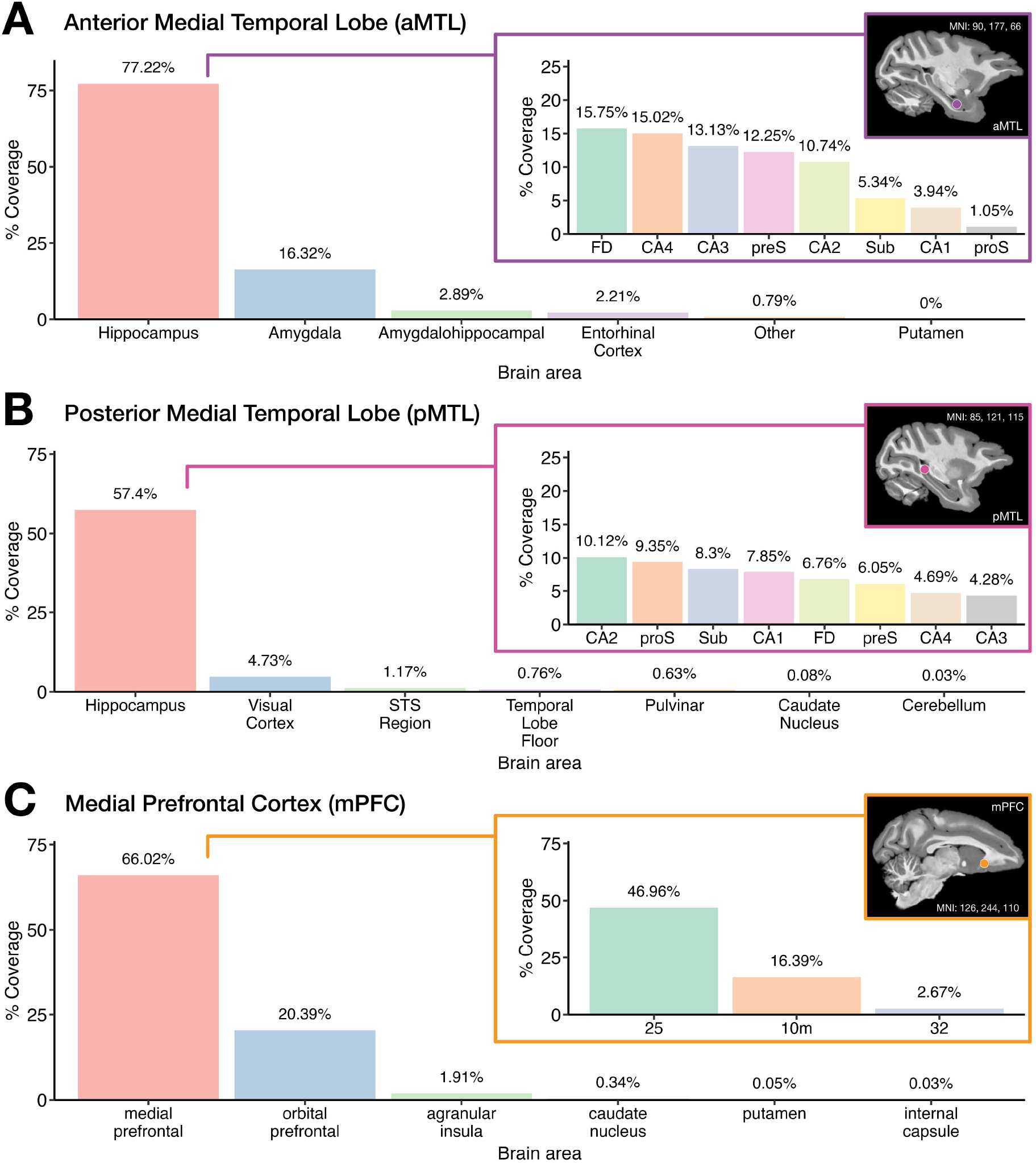
Brain regions targeted by ultrasound stimulation and their corresponding coverage in the focal ellipsoid beam modelled using the D99 atlas brain. The panels illustrate the percentage coverage of the ultrasound focal ellipsoid beam (L:10, W: 4, H: 4 mm) when targeting the three brain areas: **A)** Anterior hippocampus (aHPC), **B)** Posterior hippocampus (pHPC), and **C)** Medial prefrontal cortex (mPFC). Subplots within each panel focus on the subregions of the largest area affected. For **A-B)** this was the hippocampus, and **C)**, the medial prefrontal cortex.

## METHODS

### Experimental Model

All procedures undertaken were in accordance with the guidelines of the UK Animals (Scientific Procedures) Act of 1986, approved by Newcastle University’s Animal Welfare and Ethical Review Body and complied with the European Directive on the protection of animals used in research (2010/63/EU). All researchers conducting work with the macaques held active U.K. Home Office Personal Licenses for this work approved under an active Home Office Project License (PP8119034). All reporting follows the recommendations in the ARRIVE guidelines.

Two rhesus macaques (Macaca mulatta), one male (PL: 16 years old and 14 - 15 kg) and one female (MC: 9 years old and 6 - 7 kg) were involved in the experiment. The two macaques in this study were pair housed with other monkeys (PL with a male, and MC also with a male). The primate colony at Newcastle University is populated by approximately 40 macaques grouped in compatible social pairs or triplets. Each pair had access to a housing unit (L: 2.4 m, W: 1.4 m, H: 2.3 m) with a joining area between the two units (**Figure 1A**). The colony had a 12-hour light-dark cycle from 07:00 hr to 19:00 hr, with stable temperature (16 - 25 °C) and humidity (40 - 70 %).

All participating animals were under a Home Office Project License approved fluid control protocol to ensure sufficient motivation to conduct the behavioural task at high levels of performance, which is individually optimised for each macaque to the level of fluid control that can motivate the animal to participate in the task while ensuring sufficient fluid for their needs (at least 20 ml/kg). Testing sessions occurred during the light phase on a consistent schedule (between 9:00 hr to 13:00 hr).

### Apparatus

Behavioural testing involved the animals interacting with two touchscreens attached to two ‘rooms’ of the housing units, separated by a middle corridor (**Figure 1A**). A juice spout was located centrally below the screen and provided a fluid reward (Ribena Blackcurrant Juice, Suntory, Bristol, UK) for correctly completed trials.

### Materials

The stimuli on screen (**Figure 1B**) were photographs of real objects used in a previous context-guided decision making experiment (Slater et al., 2025). Each image measured 200 x 200 pixels, with objects presented on a neutral grey background, distinct in shape and colour to ensure rapid visual recognition and discrimination. The experiment was conducted using PsychToolbox Version-3 within MATLAB (version 2017a/2018a), running on a Windows 10 Enterprise machine connected to each touchscreen.

### Design

Prior to each testing session, the participating macaque was temporarily separated from its companion. A touchscreen was securely attached to each home-unit, and both screens were connected using a coaxial cable (Bayonet Neill-Concelman) attached to LabJack (U3, www.labjack.com) devices on each machine. This configuration ensured both touchscreens operated simultaneously on the same task.

The task is characterised as a context-dependent biconditional paired-associate task with a temporal-order component. Each context (colour cue) is associated with a consistent pair of items, defined by identity rather than screen location, and the animal must select the context-appropriate item first, then the second item, in the correct order.

An example trial is illustrated in **Figure 1C**. To initiate a trial, the macaque first touched a white circle displayed at the centre of the screen. Following a 1-second inter-stimulus interval (ISI), the stimuli appeared on the screen, prompting the animal to make a choice, ‘Response 1’ (R1). If the choice was correct, the macaques received a pre-defined volume (∼ 1.5 ml) of juice as a reward. An incorrect choice resulted in the trial being aborted without juice delivery, and a white circle reappeared to start the next trial.

Upon a correct first response, the macaque could make a second choice (‘Response 2’, R2), and if correct, a second juice reward (∼ 1.5 ml) was dispensed. After a 1-second inter-trial interval (ITI), the next trial became available. Incorrect second responses resulted in trial termination without a reward, and the white circle reappeared to initiate the next trial.

### Ultrasound Stimulation Parameters

A single-element ultrasound transducer (H115-MR) manufactured by Sonic Concepts (Bothell, WA, USA) was used in this study. Operating at a centre frequency of 250 kHz, the transducer had a radius curvature of 63.2 mm and an aperture diameter of 64 mm (**Supplementary Figure 1A, Table 1**). As such, the focal depth was fixed at 51.74 mm with the focal beam measuring (39.5 x 6.04 mm; L x W; **Supplementary Figure 1B**). To achieve a desired focal depth of 21.74 mm from the skull surface, a 30-mm coupling cone filled with degassed water was attached to the transducer housing.

**Table 1.** Transducer and Pulse Parameters.

| Parameter | Value |
| --- | --- |
| Transducer |  |
| Manufacturer (Model Number) | Sonic Concepts (H115-MR) |
| Centre Frequency | 250 kHz |
| Radius of Curvature | 63.2 mm |
| Aperture Diameter | 64 mm |
| Number of Elements | 1 |
| Matching Network |  |
| Manufacturer | Sonic Concepts (Electrical impedance matching network) |
| Pulse Timing |  |
| Pulse duration | 0.03 sec |
| Ramp Duration | 0 sec |
| Ramp Shape | Rectangular |
| Repetition Interval / Frequency | 0.1 sec / 250 kHz |
| Pulse Train Duration | 40 sec |
| Pulse Train Ramp Shape | Rectangular |

**Table 2.** Distribution of Sessions and Trials by Brain Region, Context, and Monkey.

| Phase | Monkey | sham | aMTL | pMTL | mPFC |
| --- | --- | --- | --- | --- | --- |
|  |  | (Session *n* / Trial *n*) | (Session *n* / Trial *n*) | (Session *n* / Trial *n*) | (Session *n* / Trial *n*) |
| Stable | PL | 10 / 2793 | 6 / 1565 | 6 / 1589 | 3 / 746 |
|  | MC | 12 / 710 | 3 / 554 | 9 / 349 | 4 / 113 |
| Reversal | PL | 5 / 2669 | 3 / 1626 | 3 / 1568 | 3 / 1762 |
| Unstable | PL | 9 / 2806 | 5 / 1477 | 5 / 1547 | 6 / 1682 |
|  | MC | 8 / 1033 | 7 / 1052 | 7 / 1024 | 7 / 724 |

### Drive system components

The system included a KeySight 33500B Trueform signal generator (KeySight, Santa Rosa, CA, USA) to provide the input signal required for transducer excitation. A TBS 1032B oscilloscope (Tektronix, Beaverton, OR, USA) was used for real-time signal visualisation of the signal during sonication. Signal amplification was achieved using a 75-W Model 7500 amplifier (Krohn-Hite, Brockton, MA, USA). To optimise impedance matching between the amplifier and the transducer, an electrical impedance matching network (Sonic Concepts) was included.

### Drive system settings

Sonication parameters were generated using a Windows 10 Enterprise computer and relayed to the signal generator (KeySight 33500B) via a LabJack (U3 Model). A 30-mm Perspex coupling cone filled with degassed water and sealed with a latex membrane was attached to the transducer housing to achieve the required focal length.

### Free field acoustic parameters

Free-field acoustic measurements were performed in a water bath using a 1-mm needle hydrophone (NH100, Precision Acoustics, Higher Bockhampton, DO, UK) with a measurement uncertainty of 9 %. At the focal depth of 51.74 mm, the spatial peak pressure amplitude was approximately 580 kPa, with a spatial-peak pulse-average intensity (I_sppa_) of 11.5 W / cm^2^ and a spatial-peak temporal-average *intensity (I_spta_) of 3.45 W / cm^2^*.

### Pulse timing parameters

Active ultrasound stimulation involved 30-millisecond pulses (pulse duration) delivered ultrasound every 100 milliseconds (pulse repetition interval), corresponding to a 30 % duty cycle. The total pulse train duration lasted 40 seconds (**Supplementary Figure 1C, Table 1**). The peak-to-peak voltage remained constant throughout the stimulation.

### Application of sonication

Prior to sonication, the transducer was filled with room-temperature water and sealed with a latex membrane and O-ring to eliminate air bubbles. The transducer was then calibrated to a frameless stereotaxic neuronavigation system (BrainSight Vet, Rogue Research, Montreal, CAN) using fiducial markers aligned parallel to the transducer housing. Six fiducial markers attached the animal’s head-post, along with a subject tracker, enabled calibration of the animal’s head position and orientation within BrainSight Vet. Earplugs were used throughout the sonication to protect the animal and reduce the impact of sonication noise-related confounds, although these are not expected to last beyond the sonication period (Braun et al., 2020; Kop et al., 2024).

Conductive ultrasound gel (Cutimed, Hull, Yorks, UK) was applied generously to the latex membrane. The transducer was positioned perpendicular to the target area on the scalp as guided by BrainSight and held in place for the full 40-second sonication period. Sonication was performed bilaterally in each session, with the order of hemispheres (left or right) alternated between sessions.

### Target Brain Areas

Each macaque underwent an awake T1-weighted structural magnetic resonance imaging (MRI) scan to create a template for targeting three specific brain regions: anterior hippocampus, posterior hippocampus, and the medial prefrontal cortex. Scans were acquired using a 4.7 T Bruker Vertical scanner. Six fiducial markers were affixed to the animal’s headpost to enable precise co-registration with the BrainSight neuronavigation system.

The acquired MRI scans were uploaded to the BrainSight platform where the D99 version 2.0 atlas was used as a reference to identify and map the fiducial markers and the target brain regions for sonication.

**Figure 2** illustrates the three targeted brain regions with the focal ellipsoid of the ultrasound beam overlaid, measuring approximately 10 mm in length, 4 mm in width, and 4 mm in height. The specific coordinates of the three targeted regions are as follows: medial prefrontal cortex (x: 126; y: 244; z: 110), anterior medial temporal lobe (x: 90; y:177; z: 66), and posterior medial temporal lobe (x: 85; y: 121; z: 115). This mapping allowed for precise delivery of ultrasound stimulation to the intended brain areas. **Supplementary Figure 2** highlights the coverage of the beam modelled in the D99 macaque atlas (Saleem et al., 2021) for each of the three target regions: aMTL, pMTL and mPFC.

### Behavioural Training

Habituation to the touchscreen involved two sessions per animal, allowing them to become accustomed with the touchscreens and associate touching objects onscreen with receiving a fluid reward. During this stage, objects were presented individually with reward for pressing on any of the objects, enabling the animals to familiarise themselves with the objects and background colours. Following habituation, the training procedure advanced through the following testing stages.

To minimise distractions, each animal focused on one touchscreen per session, and the touchscreens were consistently assigned to specific sides of the home-unit. Trials were counterbalanced across sessions to prevent bias or preference for a particular touchscreen, object set, or background colour.

In the initial training phase, pairs of objects were presented simultaneously in fixed spatial locations, teaching the animals the significance of object identity and sequential order within each context. Subsequent training introduced trials where objects could appear in opposite locations emphasising object identification over spatial location. Additional training steps were provided to one monkey (Monkey MC) to reinforce the importance of sequential order over spatial location.

In later training phases, incongruent objects were introduced, teaching animals to disregard specific objects irrelevant for the background colour of the screen that served as the context under which two of the specific objects were relevant in a specific order. The final phase focused on demonstrating that the spatial location of both congruent and incongruent objects was irrelevant.

### Behavioural Testing

During testing, two touchscreens were mounted to the adjacent home-units, each presenting a distinct context defined by background colour. Initially, stable contexts were used, with yellow context trials on the touchscreen to the animal’s left, and blue trials on the touchscreen on the right (**Figure 1E**). By now the two macaques understood that if the context was yellow the correct sequence order was object A followed by object B, and if the context was blue, then the sequence was object C then object D (the Stable context phase). To balance the number of trials, the active touchscreen switched after every five trials, prompting the animals to move between screens to continue the experiment.

Subsequent testing phases evaluated adaptability. While the touchscreens and contexts remained in their original positions, the object sequences were switched (**Figure 1F).** For example, in yellow context trials, object C then D became the correct sequence choice, while in blue context trials, the sequence required was object A followed by object B. This was classified as the Reversal Learning phase, which we were only able to implement with Monkey PL before all research with the macaques had to complete.

In the final testing phase, the stable assignment of contexts was removed, and yellow and blue context trials appeared randomly on either of the two touchscreens (**Figure 1G**). Novel trials introduced mid-sequence contextual changes, requiring the animals to update their responses dynamically. For example, in a trial that started with a yellow background and switched to blue mid-sequence, the animals needed to adjust their choices accordingly – initially choosing object C in the yellow context, then object B in the blue context, which was relevant to the new context. This was classified as the Unstable context phase.

## QUANTIFICATION AND STATISTICAL ANALYSIS

All statistical analyses were carried out in RStudio (R version 4.3.3, R Core Team 2023). Simulation of the ultrasound stimulation was performed in k-Plan (version 1.2.0), with further quantification of the coverage of the ultrasound ellipsoid beam being performed in MATLAB (version 2024a) and visualised in MRIcroGL (version 15.6.1).

Trial-by-trial performance (correct vs. incorrect) was modelled using bias-reduced binomial logistic regression implemented in R. The dependent variable was binary trial outcome, and the independent variable was stimulation site (sham, aMTL, pMTL, and mPFC), treated as a categorical factor. Sham trials were set as the reference category, with coefficients reflecting log-odds differences relative to sham performance. Bias-reduced logistic regression (Firth’s method) corrected for small-sample bias and significance was assessed via Wald z-tests.

Trial outcome was scored as correct only when both the first and second responses in a trial were correct. An incorrect first response terminated the trial before a second response was possible. Analyses were conducted separately for each animal, as inter-individual differences in baseline performance (sham) precluded pooling across subjects.

## AUTHOR CONTRIBUTIONS

Conceptualisation, B.S and C.P; methodology, B.S; Investigation, B.S; writing - original draft, B.S; writing - review & editing, B.S, A.E, H.C, M.K, P.D, Y.K and C.P; funding acquisition, C.P, T.G, Y.K and A.E; resources, C.P, A.E, M.K, P.D, J.S, and Y.K; project supervision, C.P, A.E, T.G and Y.K.

